# Deciphering the Cell-Specific Transcript Heterogeneity and Alternative Splicing during the Early Embryonic Development of Zebrafish

**DOI:** 10.1101/2024.09.08.611790

**Authors:** Xiumei Lin, Xue Wang, Chang Liu, Chuanyu Liu, Tao Zeng, Ziqi Yuan, Meidi Hu, Rong Xiang, Kaichen Zhao, Jie Zhou, Shichen Yang, Yang Wang, Kaifeng Meng, Hui Wang, Guangli He, Rui Zhao, Jiaheng Liu, Yunqi Huang, Jingfang Pan, Jialu Wang, Junyi Chen, Fei Guo, Yuliang Dong, Xun Xu, Daji Luo, Ying Gu, Longqi Liu, Zhiqiang Dong, Liang Chen

**Affiliations:** BGI Research, Hangzhou 310030, China; BGI Research, Shenzhen 518083, China; BGI Hangzhou CycloneSEQ Technology Co., Ltd, Hangzhou 310030, China; College of Biomedicine and Health, College of Life Science and Technology, Huazhong Agricultural University, Wuhan, Hubei 430070, China; College of Life Sciences, Wuhan University, Wuhan 430072, China; Key Laboratory of Breeding Biotechnology and Sustainable Aquaculture, Institute of Hydrobiology, Hubei Hongshan Laboratory, Chinese Academy of Sciences, Wuhan 430072, China

## Abstract

Cell fate determination during early embryonic development is a complex process modulated by gene expression. The intricate interplay of transcriptional and post-transcriptional regulation is integral to the developmental trajectory of embryogenesis, yet how RNA processing may contribute to early development programming is largely elusive. Leveraging recent technological advances in single-molecule nanopore sequencing, we developed a single-cell long-read transcriptome sequencing technology, allowing a clear view of transcript diversity during zebrafish embryogenesis, particularly spanning the periods of pre- and post-zygotic genome activation (ZGA). A closer examination of the dynamic transcript usage and potential alternative splicing revealed that abundant stage-specific transcripts with differential coding potentials are involved in distinct biological functions. Specifically, we identified two cell populations at the onset of ZGA based on isoform diversity instead of gene profiling, which followed divergent developmental trajectories toward the ectoderm and the presumptive ectoderm. These two populations of cells were characterized by divergent splicing regulations linked to differential RNA-binding proteins, including SNRPA and SFPQ. Altogether, using the single-cell long-read transcriptome sequencing strategy, our work has revealed the cell-specific transcriptome dynamics contributing to the cell fate determination during embryogenesis.

## Introduction

The early stages of embryonic development are characterized by a remarkable degree of cellular heterogeneity, with common characteristics including asymmetric cell divisions, morphogen gradients, and localized signaling centers, contributing to the determination of distinct cell fates and the formation of complex tissues and organs(*1, 2*) For example, in the developing mouse embryo, cellular heterogeneity with different activation of Hippo signal pathway is crucial for the segregation of the inner cell mass and the trophectoderm, leading to the formation of the pluripotent epiblast and the extraembryonic tissues, respectively(*3*). Similarly, in zebrafish, the differential expression of Nodal signaling components drives the formation of mesoderm and endoderm, illustrating the functional significance of cellular heterogeneity in lineage specification (*4*). On the other hand, this heterogeneity emerges through a series of highly orchestrated processes that vary across different species(*5*). In mice and humans, early asymmetric divisions can be traced back to the 2-cell stage embryo, where differential distribution of cytoplasmic determinants and localized signaling events begin to establish distinct cell fates (*1, 6–8*). In model organisms such as zebrafish, the maternal-to-zygotic transition (MZT) marks a critical period where control of development shifts from maternally deposited RNAs and proteins to zygotic gene expression, further contributing to cellular diversity(*9*). Understanding the mechanisms that drive this heterogeneity is essential for elucidating the principles of developmental biology.

The emergence of cellular heterogeneity in embryonic development is orchestrated by a complex interplay of mechanisms in across various species(*1, 10*). Previous studies have mainly focused on regulatory processes of gene expression, including signal transduction pathways, transcriptional regulation, and epigenetic modifications(*3, 10–13*) For instance, in Caenorhabditis elegans, asymmetric cell divisions are pivotal in establishing cellular heterogeneity and the anterior-posterior axis, with PAR proteins crucially partitioning cell fate determinants during early embryonic divisions(*14, 15*). The identification of morphogen gradients in Drosophila melanogaster, such as the Bicoid and Hunchback gradients, provided foundational insights into how spatial and temporal regulation of signaling pathways can drive antero-posterior axis specification(*16–18*). In Xenopus laevis, the Spemann-Mangold organizer functions as a signaling center, emitting factors like Noggin and Chordin to pattern the embryonic axis(*19–21*). In vertebrates, regulatory mechanisms become increasingly intricate. In zebrafish, a widely used model organism for studying vertebrate development, several key mechanisms contribute to early developmental heterogeneity(*12, 22*). One of the pivotal events is zygotic genome activation (ZGA), marking the transition from maternal to zygotic control of development(*23*). The dramatic reprogramming of the transcriptome during this process is one of the main drivers of cellular heterogeneity(*24, 25*). The widespread application of high-throughput single-cell sequencing technology and spatial omics technology has clarified the differentiation fate of these heterogeneous cells and the changes in gene expression along the developmental trajectory during zebrafish embryogenesis after ZGA(*26–28*). In addition to the cellular heterogeneity identified after ZGA, cell staining and microscopic imaging have shown that zebrafish embryos exhibit cellular heterogeneity and asynchronous cell division even before ZGA(*29*) However, the developmental fate and underlying mechanisms of these heterogeneous cells formed before the ZGA period remain unclear.

Transcript splicing and RNA processing play pivotal regulatory roles during embryonic development, orchestrating the precise temporal and spatial expression of transcripts necessary for cellular differentiation and tissue formation(*30*). For instance, the splicing factor Sxl (Sex-lethal) and Mbnl1 (Muscleblind-like 1) has been shown to regulate alternative splicing events crucial for sex determination, heart and skeletal muscle development(*9, 31*), demonstrating how splicing decisions can influence developmental outcomes. Furthermore, the transcript splicing processes significantly contribute to the generation of cellular heterogeneity by producing diverse transcript isoforms from a single gene, thereby expanding the functional repertoire of the genome(*32*). For example, the alternative splicing of the Dscam and Fgf8 gene produces isoforms with distinct functions in limb and nervous system development, highlighting the importance of splicing in tissue-specific transcript regulation(*33, 34*). Before the ZGA, maternal RNA processing and splicing also play a crucial role in setting the stage for subsequent developmental events. In Xenopus laevis and zebrafish, maternal mRNAs are selectively stabilized or degraded, and their splicing is tightly regulated to ensure proper embryonic development(*35, 36*). Additionally, the gene *YBX1*, associated with mRNA splicing and stability in mice, is crucial for embryonic development by regulating the alternative splicing of maternal mRNA and activating early zygotic genes(*37*). In vitro experiments with Pladienolide B (PlaB) demonstrated that inhibiting spliceosome function can re-induce pluripotent cells into totipotent stem cells, underscoring the necessity of normal alternative splicing for the transition from totipotency to pluripotency in early embryos(*38*). Therefore, systematically investigating alternative splicing events and transcript isoforms of maternal and zygotic genes using a model animal like zebrafish offers a novel perspective to understanding the mechanisms behind early embryonic cell heterogeneity, providing critical regulatory insights previously unattainable at the gene expression level.

The advent of long-read sequencing technologies has revolutionized our ability to study transcriptomic complexity. Unlike short-read sequencing, which often struggles with accurately reconstructing full-length transcripts, long-read sequencing provides the capability to sequence entire RNA molecules in a single read(*39*). This allows for the precise identification of transcript isoforms, including those generated by alternative splicing, and provides a more comprehensive view of the transcriptome. Advances in long-read sequencing technologies have enabled the comprehensive profiling of full-length transcript isoforms, revealing the complexity of splicing events and their regulatory roles in development(*40*). Recently, researchers have leveraged long-read sequencing technologies, such as PacBio’s Single Molecule, Real-Time (SMRT) sequencing and Oxford Nanopore’s platforms, to develop single-cell full-length transcriptomes(*41–43*). These advancements have extended beyond transcriptomics to include epigenomics, 3D genome organization, and other multi-omics analyses, offering deeper insights at single-cell resolution into the molecular mechanisms driving critical life processes such as nervous system function and early embryonic development(*44–46*). These technologies have enabled the exploration of cellular heterogeneity, revealing intricate details about cell-specific transcriptomes and the dynamic changes that occur during development.

Despite these advancements, the lack of comprehensive single-cell long-read transcriptomic studies on embryogenesis limits our ability to fully understand the complexity of transcript usage and alternative splicing, as well as its impact on cell fate determination. Here, we adopted the newly developed single-molecule nanopore sequencing platform (CycloneSEQ) (*47*)to establish a single-cell long-read transcriptome sequencing technology and dissect the global transcript heterogeneity landscapes based on comprehensive full-length transcript information during the critical window before and after ZGA of zebrafish embryo development. Our study provides important data resources and a robust and comprehensive full-length transcript data analysis methodology for researching the usage diversity of transcript isoforms and alternative splicing events during vertebrate embryogenesis.

## Results

### Full-length transcriptome diversity in zebrafish early embryonic development

Precise single-cell barcode demultiplexing is challenging and error-prone for long-read sequencing, due to the lower sequencing accuracy compared to the next-generation sequencing (NGS) platforms. To address this issue, we prepared a mix of cell suspension containing an equal amount of human (HEK293T) and mouse (NIH-3T3) cells for scRNA-seq library construction using the DNBelab C4 Droplet platform. Subsequently, we conducted the short-read sequencing by DNBSEQ and long-read sequencing by CycloneSEQ in parallel(*47*). By aligning to the valid cell barcode whitelist generated from short-read sequencing, we successfully demultiplexed single-cell barcodes for CycloneSEQ reads with high sequence consistency for analysis. (fig. S1A) (see Methods). To assess the accuracy of barcode demultiplexing, we conducted mixed-cell analyses on two datasets, revealing comparable mixing rates between the two replicates (fig. S1B). Additionally, we compared single-cell transcriptomes generated by CycloneSEQ with those from standard DNBSEQ and noted a high degree of concordance (fig. S1C), confirming the reliability of the barcoding detection approach. Consequently, we established the single-cell long-read sequencing technology by integrating short-read and long-read data to construct a more detailed information matrix that encompasses transcript expression and usage differences, thereby surpassing the limitations of traditional gene-level analytical approaches.

Zebrafish provide a vital model for embryogenesis research, enabling detailed investigation of molecular mechanisms from fertilization to lineage development due to their external and rapid early development(*48, 49*). We systematically investigated the dynamic changes in RNA processing and transcriptome diversity during zebrafish embryogenesis throughout MZT using long-read sequencing. Zebrafish embryos from various developmental stages including 128-cell (2.25 hours post-fertilization (hpf)), 512-cell (2.75 hpf), 1k-cell (3 hpf), Dome (4.3 hpf), and 50%-epiboly (5.25 hpf), were harvested for full-length transcriptome analysis (Fig.1A). We filtered for transcripts with an expression level greater than 5 (umi > 5) detected in at least three cells, identifying a total of 27,717 transcripts. Among these, 18,360 (66.24%) were full splicing isoforms (FSM), while 7,051 (25.44%) represented novel transcripts (Novel In Catalog (NIC) = 3.50%, Novel Not In Catalog (NNIC) = 21.92%) (fig. S1D). Additionally, the transcription start sites (TSS) of the representative novel transcripts were supported by snATAC-seq peaks(*50*) (fig. S1E). Despite no discernible variation in length across transcript types, the novel transcripts exhibited significantly reduced expression levels compared to annotated transcripts, consistent with previous studies(*51*) (fig. S1, F and G). Both annotated and novel transcripts predominantly displayed protein-coding potential and maintained relative consistency across various developmental stages (fig. S1, H and I). The majority of genes corresponded to only one or two transcripts (fig. S1J), and a concordance was observed between genes and transcripts at each developmental stage (fig. S2, A and B).

**Fig. 1.**
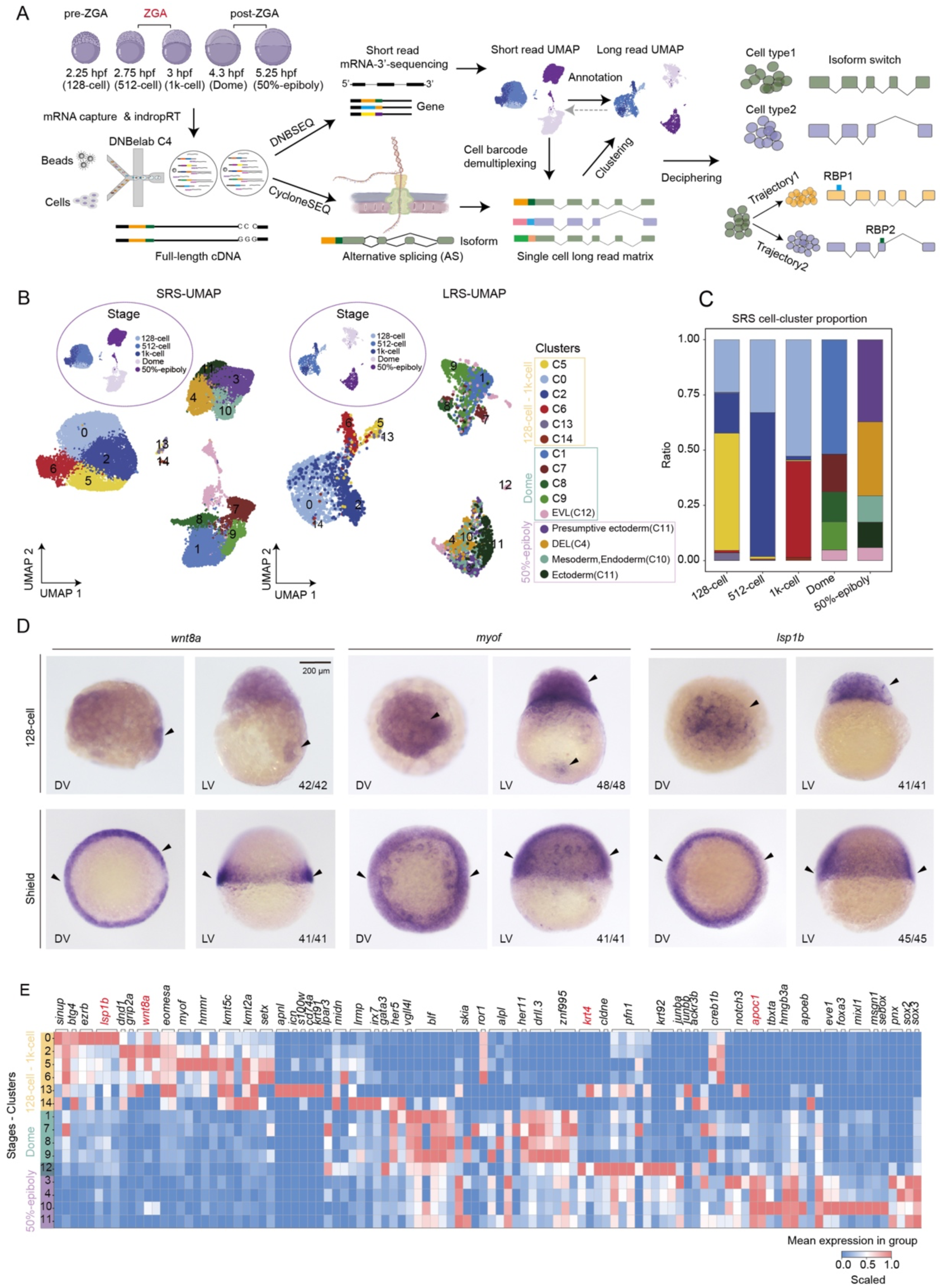
Full-length transcriptome diversity in zebrafish early embryogenesis. (A) Schematic illustrating transcriptomic diversity and alternative splicing in zebrafish embryogenesis, resolved with single-cell DNBSEQ and CycloneSEQ technologies. (B) UMAP plots illustrating cell clusters and stages: 28,468 cells detected by short read sequencing (left) and 3,851 metacells detected by long read sequencing (right). Colored by clusters and the developmental stages. (C) A bar chart shows cluster proportions across five developmental stages in short-read sequencing. The color legend of is the same as that in Fig.1B. (D) In situ imaging of *wnt8a*, *myof* and *lsp1b* at the 128-cell and shield stages. Scale bar, 0.2 mm. (E) The heatmap displays the expression of cluster specific genes with various isoforms by long-read sequencing, though isoform details are omitted from the heatmap.

Using short-read sequencing data, we integrated and clustered a total of 28,468 cells across five developmental time points, identifying 15 distinct cell clusters (Fig. 1B). Notably, prior to ZGA (at the 128-cell, 512-cell, and 1k-cell stages), there was considerable overlap among cell clusters (Fig. 1, B and C). These cell types were distinguished by specific marker genes: C0 exhibited high expression of *lsp1b*, C5 highly expressed *myof*, and C2 primarily expressed *grip2a* and *wnt8a* (fig. S2C). Unlike the well-documented *grip2a* and *wnt8a* functioning in dorsal-ventral axis formation in zebrafish(*12, 52*), the contributions of *lsp1b* and *myof* to early body axis patterning are yet to be defined. We therefore performed the *in situ* hybridization assay and observed a similar spatial expression pattern for *lsp1b* and *wnt8a* and a pronounced expression of *myof* in the ventral region (Fig. 1D). These findings hint at possible involvement of *lsp1b* and *myof* in dorsal-ventral axis development, while further research is needed to clarify their roles. However, prior to the 50%-epiboly stage, assigning definitive cellular identities remained challenging despite the presence of differential expression patterns. At 50%-epiboly, distinct cell types became discernible: C11 ectoderm was characterized by high levels of *sox2* and *sox3*, C10 mesoderm and endoderm by *msgn1* and *foxa3*, and the enveloping layer (EVL) by *krt4* and *cldne*. *Notch3*, crucial for the self-renewal and differentiation of cytotrophoblast progenitors in early human placental development(*53*), was prominent in C3, where *sox2* and *sox3* were expressed at lower levels, suggesting a presumptive ectoderm status. C4, lacking distinctive markers, was classified as deep cell layer (DEL) (Fig. 1B and fig. S2C).

Contrasting the distinct cell clustering result based on short-read data, the cell clusters with the long-read data becomes unclear due to the data sparsity (3 k reads/cell) (fig. S2, D and E). To resolve this issue, we adopted the SEACell strategy(*54*). We assigned the cell cluster information from short-read data to long-read data based on cell IDs and grouped cells into metacells within each cluster according to expression similarity (see Methods), yielding 3,851 metacells with mean gene expression exceeding 2000 (fig. S2F). Based on this matrix, we conducted cell clustering for long-read data and integrated the cluster information from short-read data (Fig.1B), revealing expression consistency between the two datasets (fig. S2, G and H). Our analysis of cell type-specific markers and their corresponding transcript information revealed that genes such as *wnt8a*, *krt4*, *apoc1* and *lsp1b* exhibit multiple transcript variants across cell types (Fig. 1E and fig. S2, I and J). In summary, we have developed a single-cell long-read sequencing technology integrated with the SEACell strategy, effectively addressing the sparsity challenges associated with long-read data and providing a comprehensive, transcriptome-wide resource throughout the MZT.

### Global changes in transcriptome usage during zebrafish MZT

Transcript usage preference analysis provides a more nuanced understanding of gene functional changes throughout development than merely assessing gene expression levels(*55*). Although MZT has been extensively studied at the level of transcription and epigenetic regulation(*56*), the dynamic alterations in transcript usage during this critical developmental period are merely explored. To identify specific transcript usage at each stage, we scrutinized isoform usage specific to each stage (see Methods). Our results revealed that the predominant transcripts at the 128-cell stage were primarily associated with genes involved in cell cycle, ribosome activity, and RNA metabolism, while transcripts predominantly used during the 512-cell and 1K-cell stages were linked to genes related to cell cycle transitions and gastrulation (Fig. 2A). Notably, the cell division-related gene *ccnd1* displayed different transcript variants with stage-specific usage patterns, suggesting that *ccnd1* has a dominant transcript at each developmental stages and potentially performs different functions (Fig. 2B).

**Fig. 2.**
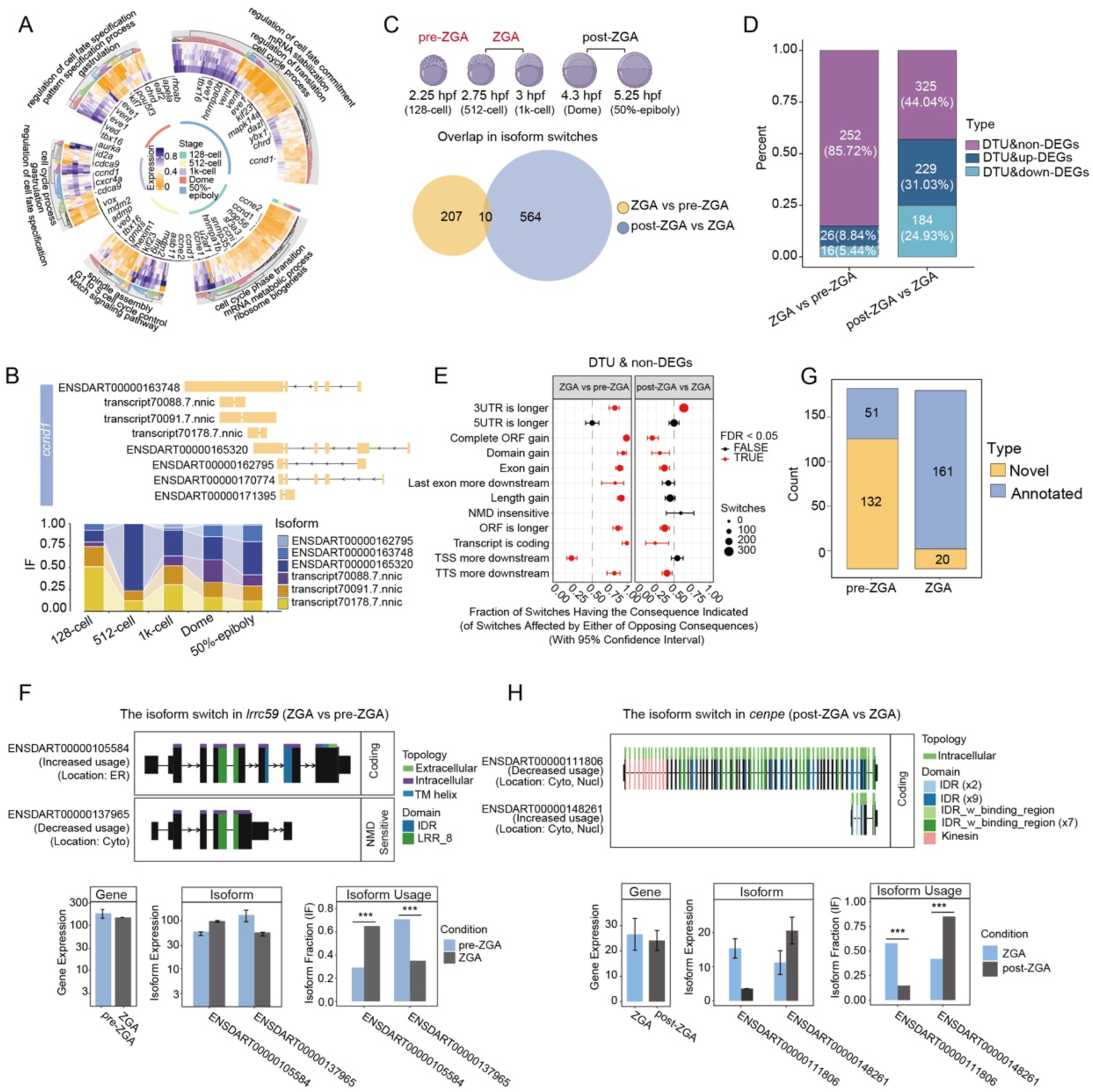
Global changes in transcriptome usage during zebrafish MZT. (A) A circle heatmap illustrates differential transcript usage (DTU) patterns across developmental stages in long-read sequencing data. dIF > 0.1 and adjusted p-value < 0.05. (B) A schematic shows *ccnd1* transcript structure with a stacked bar chart detailing isoform fraction (IF) values at various stages. (C) The Venn diagram illustrates isoform switch events, comparing ZGA to pre-ZGA and post-ZGA to ZGA. (D) The stacked bar chart shows DTU and DGE between pre-ZGA vs ZGA, and ZGA vs post-ZGA. DTU is defined by dIF > 0.1 and p < 0.05, while DGE is categorized as upregulated (log2FC > 0.5, p < 0.05), downregulated (log2FC < -0.5, p < 0.05), or stable otherwise. (E) The illustration depicts functional consequences of isoform switching in non-DGE categories, comparing ZGA to pre-ZGA and post-ZGA to ZGA. (F) Isoform switching of the *lrrc59* gene was observed between pre-ZGA and ZGA stages. A bar chart illustrates the gene expression, isoform expression, and isoform usage. (G)The stacked bar chart displays the counts of annotated and novel transcripts, exhibiting isoform switching in non-DEG categories between pre-ZGA and ZGA stages. (H) Isoform switching of the *cenpe* gene observed between ZGA and post-ZGA stages. A bar chart shows the gene expression, isoform expression, and isoform usage.

To better understand transcript usage dynamics around ZGA, we categorized five developmental stages into three phases: pre-ZGA (128-cell stage), ZGA (512-cell and 1K-cell stages), and post-ZGA (Dome and 50% epiboly stages). Our findings indicated relatively few changes in transcript usage during ZGA, with more significant shifts occurring afterward (Fig. 2C). Initially, at ZGA, differential usage was prevalent in non-differentially expressed genes (non-DEGs), but this trend reversed post-ZGA, where over half of the changes involved differentially expressed genes (DEGs) (Fig. 2D). Moreover, transcript usage in non-DEGs during ZGA was characterized by a preference for more complete transcripts, featuring longer 3’-UTR (approximately 80%), completed open reading frame (ORF) gain (approximately 90%), and exon gain (approximately 87.5%), which are crucial for cell cycle regulation, transcription, and mRNA processing (Fig. 2E and fig. S3A). For example, *lrrc59*, key to mRNA translation and cell cycle regulation, underwent a significant isoform switch during ZGA. Specifically, Nonsense-Mediated Decay (NMD)-sensitive transcripts ENSDART00000137965 were prevalent before ZGA but diminished thereafter, coinciding with an upregulation of complete transcripts ENSDART00000137965 (Fig. 2F). Additionally, we observed a notable decrease in the expression of novel transcripts that were prevalent in the pre-ZGA phase, coinciding with a rise of annotated transcripts during ZGA (Fig. 2G and fig. S3B). This transition from novel- to annotated-transcripts indicated the significant shifts in functions related to cell cycling and RNA metabolism during ZGA activation.

Contrasting with the ZGA phase, we observed a notable rise in novel transcripts among those with differential usage for both non-DEGs and DEGs post-ZGA (fig. S3, C to F). This trend was associated with a preference for shorter ORF transcripts in the downregulated genes post-ZGA (fig. S3G), particularly in cell cycle and DNA damage response genes (fig. S3H). For instance, *cenpe*, essential for zebrafish zygotic cell division(*57*), preferred to be downregulated. Moreover, the transcripts that encoded shorter ORFs with fewer functional domains significantly increased in the post-ZGA phase (Fig. 2H), reflecting the gene regulation at both transcription and post-transcription level. Conversely, in the upregulated genes, most transcripts exhibited longer 3’ UTRs in the post-ZGA phase (fig. S3G). The extension of 3’ UTRs has been reported to regulate mRNA and protein translation efficiency, degradation, and subcellular localization through microRNAs (miRNAs) and RNA-binding proteins (RBPs) (*58*). The functions of these genes with longer 3’UTRs are mainly central to RNA splicing and processing (fig. S3H), indicating an increase in translational demand to support developmental progression post-ZGA. In summary, our single-cell long-read data analysis of zebrafish embryogenesis underscores a close association of differential transcript selection and gene expression with developmental transitions before and after ZGA.

### Global changes in alternative splicing during zebrafish MZT

Alternative splicing is an important post-transcriptional regulatory mechanism in eukaryotes that plays a critical role in cell differentiation, organ development, and tissue homeostasis maintenance(*59*). To elucidate the global changes in alternative splicing, as well as the potential interplay between gene expression and alternative splicing pre- and post-ZGA at the stage specific manner in zebrafish embryos, we employed Percent Spliced In (PSI) metric, gene expression level and exon inclusion level in our analysis (Fig. 3A) (see Methods). In contrast to the limited differential transcript usage between pre-ZGA and ZGA phases, we detected a comparable number of differential splicing events for the two comparisons (367 at ZGA vs Pre-ZGA and 436 at post-ZGA vs ZGA) (Fig. 3B). In ZGA compared to pre-ZGA, most splicing differences occurred in non-DEGs. Whereas in post-ZGA phase compared to ZGA, 66.50% of differential splicing events were associated with DEGs (Fig. 3C). To clarify the relationship between splicing events and DEGs, we selected genes with differential exon usage among the DEGs and categorized them into upregulated and downregulated groups (Fig. 3, D and E). Upregulated genes, such as *lin28a* and *zeb1a*, showed a higher incidence of exon inclusion events, whereas for genes encoding RNA-binding proteins, like *elval1a* and *hnrnpk,* a gradual transition from inclusion to exclusion was observed over time (Fig. 3D). Similarly, downregulated genes involved in cell cycle regulation, including *ccni, cdc20*, *anln* and *kif2c*, also displayed variations in exon usage (Fig. 3E).

**Fig. 3.**
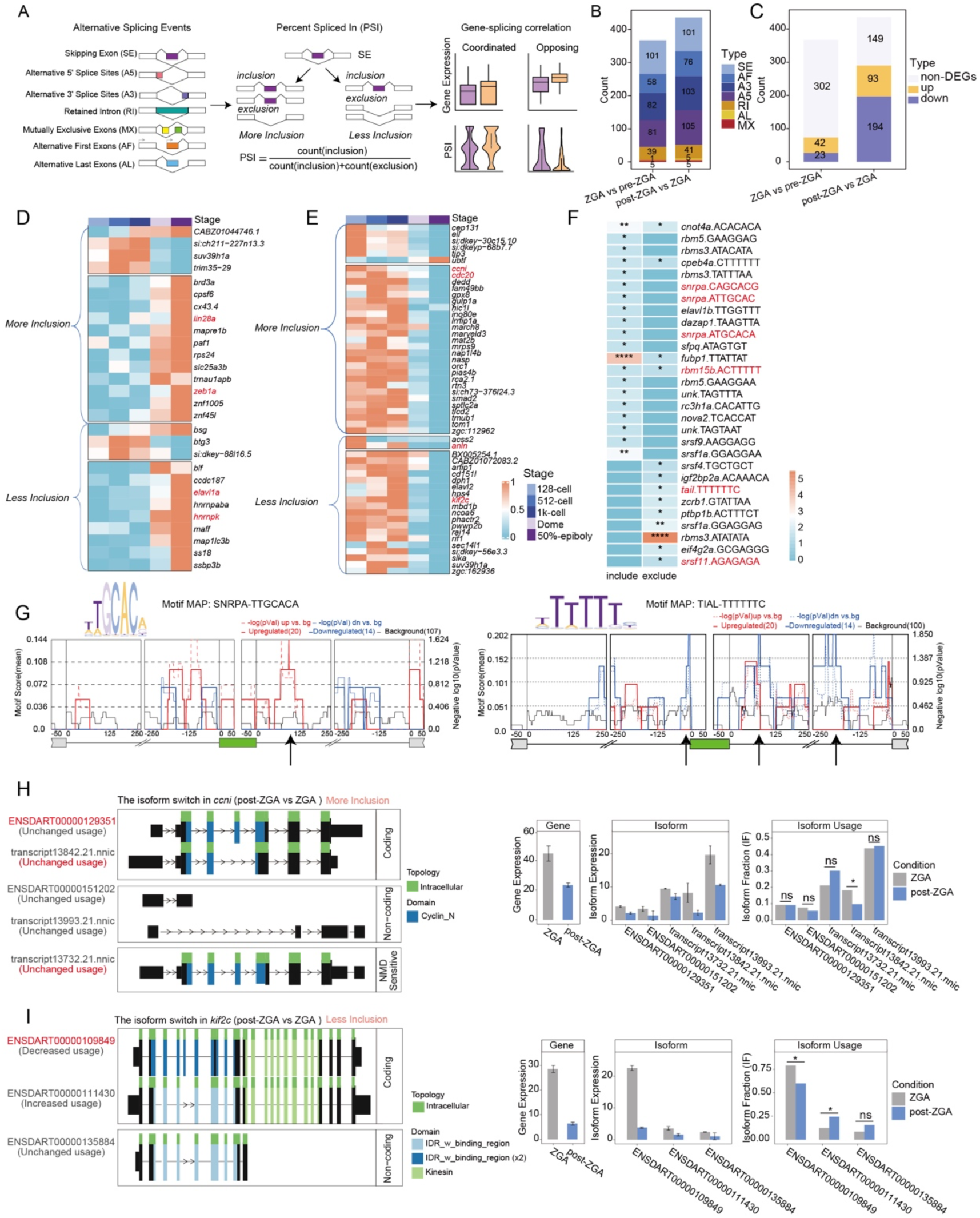
Global changes in alternative splicing during zebrafish MZT. (A) The diagram illustrates splicing event patterns and PSI calculation. (B) The stacked bar chart displays the proportion of differential splicing events (DPSI) between ZGA vs pre-ZGA and post-ZGA vs ZGA. |DPSI| > 0.05 and p-value < 0.05. (C) The stacked bar chart visualizes the relationship between the differential splicing events and DEGs, upregulation gene: adjusted p-value < 0.05 and log fold change > 0.5; downregulation gene: adjusted p-value < 0.05 and log fold change < -0.5. (D) The heatmap visualizes upregulated genes across developmental stages that correspond to DPSI in exon skipping events. DPSI > 0.05 and adjusted p-value < 0.05. (E) The heatmap visualizes downregulated genes across developmental stages that correspond to DPSI in exon skipping events. DPSI < 0.05 and p-value < 0.05. (F) The heatmap displays the enrichment results of RNA-binding protein (RBP) motifs corresponding to DPSI in exon skipping event across time points. |DPSI| > 0.05 and p-value < 0.05. (G) The enrichment results for the SNRPA and TIAL1 RNA-binding proteins (RBPs) are depicted, with arrows indicating the binding positions of these RBPs. (H-I) The switch plot presents results for the *cnni* (H) and *kif2c* (I) between ZGA and post-ZGA. A bar chart illustrates gene expression, isoform expression and isoform usage.

RNA-binding proteins (RBPs) are key mediators of post-transcriptional RNA regulation(*60, 61*). To better understand the role of RBPs in alternative splicing throughout zebrafish embryogenesis, we performed the RBP motif prediction. First, we grouped these differential splicing events based on the occurrence of more inclusion or exclusion events during MZT. By employing rMAPS, we enriched for RBP motifs, thereby identifying 27 RBPs potentially involved in the regulation of alternative splicing (Fig. 3F) (see Methods). To find RBPs more closely associated with these splicing events, we selected 26 RBPs that showed gradually increasing expression over time (fig. S3I). Among these, we identified overlaps with candidate RBPs coding genes such as *snrpa*, *tial*, *rbm15b* and *srsf11* (Fig. 3F and fig. S3I). Notably, SNRPA binds to introns flanking target exons, facilitating the retention of those exons (Fig. 3G, left). In contrast, the RBP TIAL, which also increased over time, was primarily enriched in splicing events involving exclusion, suggesting a role in promoting exon skipping (Fig. 3G, right). To analyze which genes these RBPs might influence, we selected genes with differential splicing events across developmental stages. Among the downregulated genes, *ccni* exhibited increased usage of exon-included transcripts ENSDART00000129351 and transcript13732.21.nnic post-ZGA, while *kif2c* favored exon-skipped transcripts, showing a significant decrease in the expression of the exon-included transcript ENSDART00000109849 (Fig. 3, H and I). Taken together, our study provides insights into the nuanced dynamics of alternative splicing during zebrafish embryogenesis, identifying RNA-binding proteins (RBPs) such as SNRPA and TIAL in the regulation of exon inclusion and exclusion around the ZGA. Additionally, we observed stage-specific splicing variations in genes like *ccni* and *kif2c*.

### Isoform diversity uncovering finer subpopulations

Previous research demonstrates that the transcript diversity analysis may uncover intermediate transitional cell types(*62*). To evaluate whether the transcript expression profiles infer more delicate state differences between cells than gene expression, we performed the entropy analysis to estimate the degree of expression heterogeneity for genes or transcripts within each cluster (see Methods). The results indicate that the gene expression profile of cells within a cluster is highly homogeneous for most cell clusters (rogue > 0.9), yet considerable intra-cluster heterogeneity at the transcript level (Fig. 4A). Therefore, we performed sub-clustering based on transcript diversity until the entropy values exceeded 0.9 for the majority of cells in a sub-cluster (fig. S4A) and identified a total of 24 sub-clusters. Among these clusters based on DEGs, further refinement was possible by leveraging transcript heterogeneity (Fig. 4B). The cluster C0 exhibited the highest heterogeneity and could be further divided into four major sub-clusters (Fig. 4C). Moreover, we conducted an in-depth analysis to explore isoforms and splicing diversity at the gene level across various cell clusters. The results revealed a modest increase in transcript diversity within groups C0 and C2, marked by a slightly higher proportion of genes with three or more isoforms (fig. S4B), while no significant variations were observed in the composition and proportion of splicing events (fig. S4C).

**Fig. 4.**
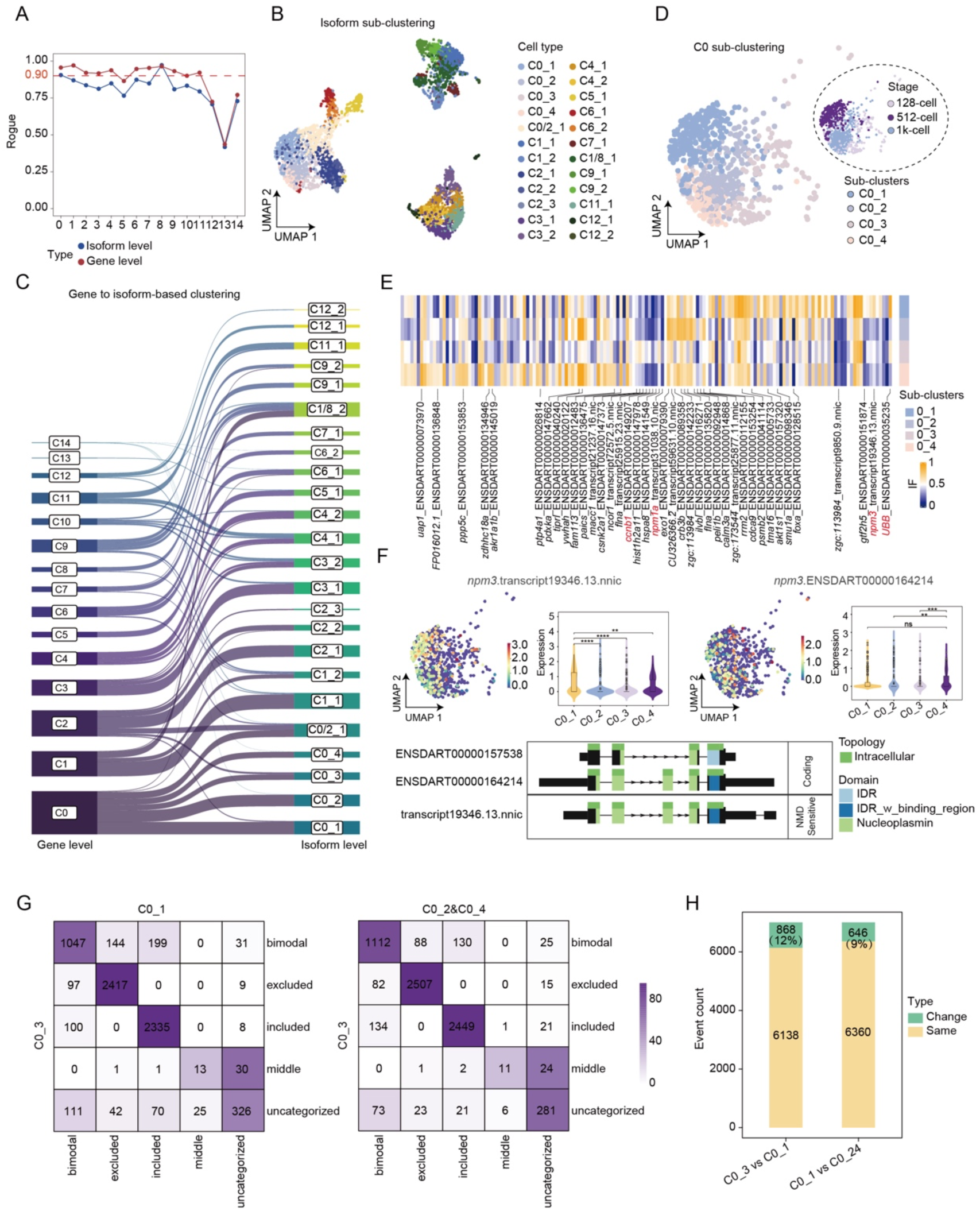
Isoform diversity uncovering finer subpopulations. (A) A line chart displays cellular heterogeneity at gene and transcript levels in long read data, as assessed by ROGUE. (B) A UMAP plot illustrates detailed sub-clustering at the transcript level from long read data. (C) River plot showing mapping of cells from gene-based (left) to isoform-based (right) clustering. Each line represents a single cell. (D) A UMAP plot displays the sub-cluster C0 at the transcript level from long read data. (E) Heatmap showing the isoform faction with DTUs between sub-cluster in C0. dIF> 0.1 and adjusted p-value < 0.05. (F) Violin and UMAP plots (top) display the expression of *npm3* transcripts within the C0 sub-cluster. The switch plot (bottom) presents the results for *npm3* transcript isoform switching. (G) Stacked bar chart illustrating modal switching events between C0_3 and C0_1, and between C0_3 and C0_2&C0_4. (H) Stacked bar chart illustrating modal switching events between C0_3 to C0_1 and C0_3 to C0_2 & C0_4.

Next, we conducted a detailed analysis of cluster C0, identifying four sub-clusters that closely correspond to three developmental stages: 128-cell (mapped to C0_3), 512-cell (mapped to C0_1) and 1k-cell (mapped to C0_2 and C0_4). This suggested a temporal progression of cellular sub-clusters defined by transcriptomic diversity, despite minimal variations in overall gene expression (Fig. 4D). Our analysis of transcript usage preferences revealed a nuanced pattern of transcript selectivity among these sub-clusters. Notably, the sub-cluster C0_1 specifically favored transcripts ENSDART00000035235 (one transcript of *ubb*) and transcript19346.13.nnic (one transcript of *npm3*), which are implicated in proteasomal degradation(*63*) and chromatin remodeling(*64*), respectively (Fig. 4E). Sub-cluster C0_3 preferentially expressed transcripts, including transcript31038.10.nic from *npm1a*, a gene known for its roles in ribosome biogenesis, centrosome duplication, and hematopoiesis(*65*), as well as ENSDART00000149207 from *ccnb1*, which is involved in cell cycle regulation.

To identify sub-cluster-specific functions of differentially distributed transcripts, we performed structural prediction analysis. Notably, the predominant isoforms of *npm3* (transcript19346.13.nnic) and *npm1a* (transcript31038.10.nic) in C0_1 and C0_3 were tended to undergo NMD and predicted to be non-coding, respectively, which are in accordance with the downregulated levels at latter (Fig. 4F and fig. S4D). Conversely, *ubb (*ENSDART00000063357) and *ccnb1*(ENSDART00000035235) tend to utilize transcripts with more complete structural domains in C0_1, indicating their roles in cell cycle progression and proteasomal degradation (fig. S4, E and F). This underscores the importance of transcript usage in modulating gene activity in a stage-specific manner. In conclusion, focusing solely on gene expression levels can overlook the functional roles of specific transcripts. Our analysis underscores that transcript usage is pivotal for cellular function and development, offering a refined tool for deciphering cellular heterogeneity.

To further explore the relationship between alternative splicing and differential transcript usage among cell sub-clusters, we employed a splicing modalities analysis, which assesses changes in splicing levels at the single-cell level(*66*). From sub-cluster C0_3 (128-cell stage) to C0_1 (512-cell stage), 12% of splicing forms underwent modal transitions, primarily shifting from bimodal to inclusion or exclusion patterns. However, the change in these splicing modalities from C0_1 to C0_2/C0_4 (1k-cell stage) showed a slight decrease, at only 9% (Fig. 4, G and H). In addition to transitions from multimodal to inclusion or exclusion patterns, there was also a notable increase in shift of inclusion towards bimodal splicing patterns from C0_1 to C0_2/C0_4 (Fig. 4G), which may reflect an upregulation of splicing activity associated with the ZGA.

### Isoform subgrouping associated with distinct developmental trajectories

The majority of cell trajectory analyses are based on gene expression data, leaving the influence of transcript diversity on developmental trajectories an open question. To this end, we utilized fine-grained transcript sub-clusters to estimate the potential developmental trajectories using the KNN algorithm(*67*), leading to the construction of a transcript-based trajectory (see Method). We identified that the transcript sub-clusters C2_1 and C2_2 from the 512-cell stage C2 diverge along two distinct developmental trajectories, giving rise to cell types Presumptive ectoderm (C3_1: trajectory 2) and Ectoderm (C11_1: trajectory 1) at the 50%-epiboly stage (Fig. 5A and fig. S5A), each characterized by different expression profiles, such as *sox3* in the Ectoderm and *celsr1a*(*68*) in the Presumptive Ectoderm (Fig. 5B). To ascertain the role of alternative splicing in shaping distinct developmental trajectories, we analyzed the differential splicing events between trajectories 1 and 2. Our comparison revealed a higher incidence of splicing events in trajectory 1, predominantly characterized by skipped exons (SE) and alternative 3’ splice site (A3) events (Fig. 5C). The genes undergoing differential splicing between the trajectories highlighted that trajectory 1 was predominantly associated with RNA processing, translation, and convergent extension (fig. S5B). In contrast, trajectory 2 was primarily linked to RNA metabolic processing and polycomb repressive complex (fig. S5B), with genes such as *bmi1a* and *rbbp4* involved in regulating gene expression and maintaining cellular pluripotency(*69, 70*).

**Fig. 5.**
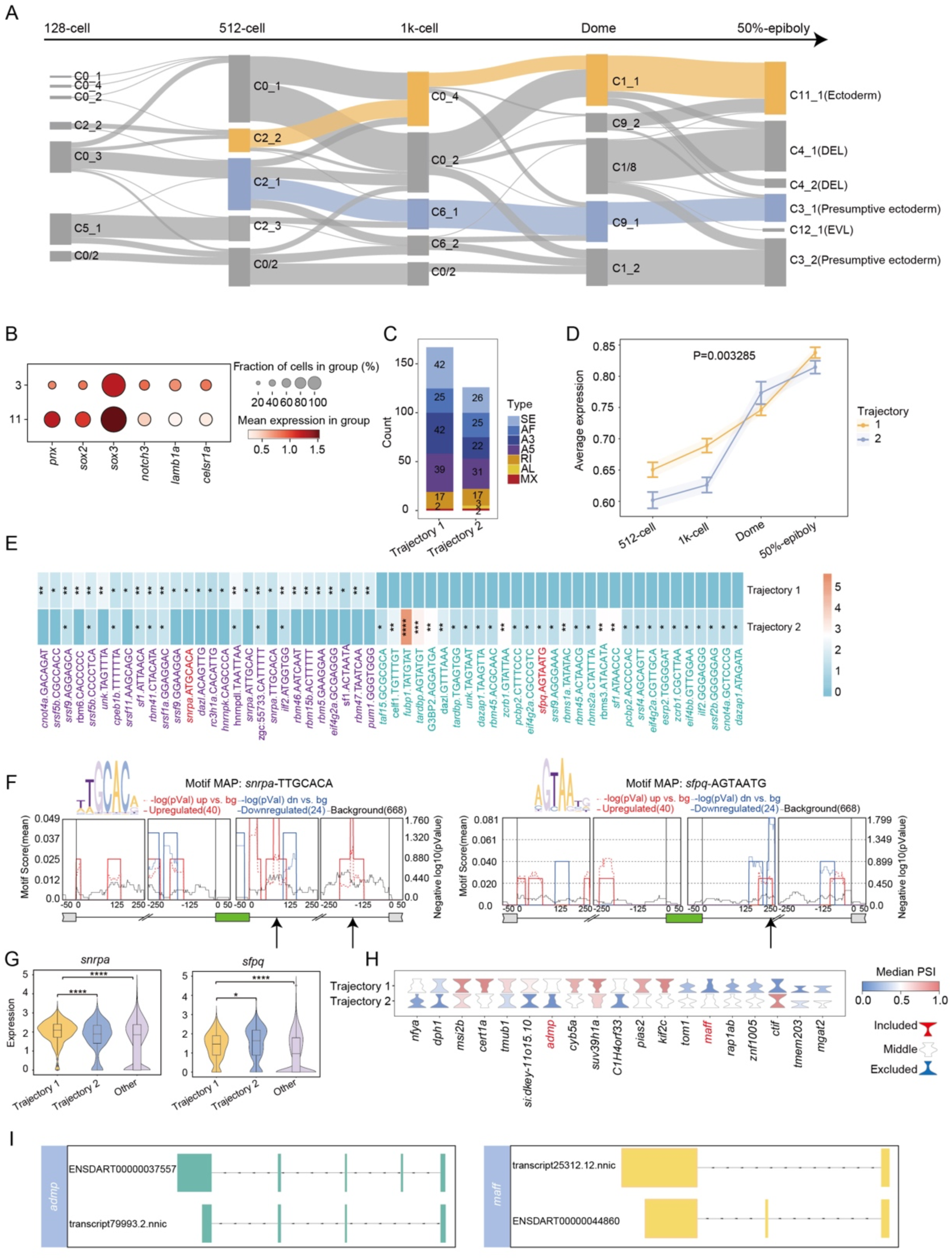
Isoform subgrouping associated with distinct developmental trajectories. (A) A Sankey diagram visualizes the developmental trajectory of zebrafish embryos, incorporating isoform sub-clusters identified in long read data. Two developmental branches, leading to the ectoderm and presumptive ectoderm, are highlighted with distinct colors. (B) A dotted heatmap illustrating the marker gene expression profiles for presumptive ectoderm (C3) and ectoderm (C11). (C) The stacked bar chart displays the proportion of differential splicing events between two trajectories. ΔPSI > 0.1 and p < 0.05. (D) Line chart of RBP expression profiles from long read sequencing across time points in two trajectories. (E) The heatmap displays the enrichment results of RNA-binding protein (RBP) motifs corresponding to differential splicing events (DPSI) associated with exon skipping event between two trajectories. P < 0.05 and ΔPSI > 0.1. (F) The enrichment results for the SNRPA and SFPQ RNA-binding proteins (RBPs) are depicted, with arrows indicating the binding positions of these RBPs. (G) Violin plot depicting the expression patterns of *snrpa* and *sfpq* across two specified trajectories in comparison to other trajectories. (H) Violin plot showcasing genes with differential exon usage between two trajectories. ΔPSI > 0.3 and p < 0.05. (I) Transcript structure diagram of *admp* and *maff*.

We noted pronounced disparities in RBP expression between two trajectories: trajectory 1 displayed elevated RBP levels before ZGA (1k-cell stage), yet these differences waned post-ZGA, while the expression of the housekeeping gene actb1 remained constant (Fig. 5D and fig. S5C). To elucidate the RBPs driving these splicing variations, we conducted motif predictions for RBPs on differentially exon skipping events, revealing 28 motifs preferentially associated with trajectory 1 and 30 with trajectory 2 (Fig. 5E). The expression profiles of these RBPs mirrored this trend, showing marked divergence pre-ZGA (1k-cell), followed by a convergence (fig. S5D). Among these RBPs, SNRPA is mainly enriched in trajectory 1 and SFPQ in trajectory 2, participating in the regulation of alternative splicing via interactions with intronic regions of target genes (Fig. 5, E and F). Additionally, SFPQ has been reported to play a role in the development of the zebrafish nervous system(*71*). In addition, both splicing factors exhibited significant trajectory-specific expression compared to others (Fig. 5G). Upon closer examination of genes exhibiting differential exon usage between the trajectories, we observed distinct splicing patterns (Fig. 5H), even when the overall gene expression levels were comparable (fig. S5E). For example, *admp*, which is expressed in the anterior neuroectoderm and mesoderm during gastrulation(*72*), showed different transcript usage between the trajectories: trajectory 1 favored exon inclusion, whereas trajectory 2 had higher levels of exon-skipped transcripts (Fig. 5I and fig. S5F). In conclusion, transcript sub-clustering revealed two developmental branches with distinct differentiation profiles. The early-stage RBP and splicing variations between these branches highlight the potential of alternative splicing to significantly influence cell fate determination at the onset of cellular commitment.

## Discussion

Comprehensively employing the full-length transcript information through the single-cell long-read sequencing technology, our study elucidated, for the first time, the intricate isoform diversity, dynamic isoform usage, and alternative splicing in zebrafish embryos during MZT. Meanwhile, we developed a robust and comprehensive data analysis pipeline that encompasses long-read barcode demultiplexing, differential transcript usage, and alternative splicing analysis at single-cell resolution. This workflow is designed to reveal transcript heterogeneity and differential splicing among cell clusters, shedding light on the complexities of transcriptome diversity and the underlying regulatory mechanisms within the cellular. Our findings underscore the essential roles of isoform dynamics and alternative splicing in early development, complementing gene expression regulation. Simultaneously, the full-length transcript information enables us to delve deeper into the transcript usage switch over development, predict the biological functions of transcripts specific to various developmental stages, and infer the RBPs that regulate the corresponding splicing events. Moreover, at the onset of ZGA, we identified two distinct cell subpopulations with isoform diversities and divergent developmental pathways that converge on the ectoderm, each characterized by unique differentiation sequences. Notably, gene expression alone was insufficient to distinguish these diverse cell subclusters. Therefore, single-cell long-read sequencing technology has provided unprecedented insights into the complexity of transcriptome regulation during early embryonic development, underscoring the critical role of isoform diversity and alternative splicing in shaping cellular heterogeneity and developmental trajectories.

Around the critical developmental period of ZGA, we investigated developmental stage- and cell-specific isoforms. Yet, the functional significance of the numerous low-expression transcripts present remains unclear. In our analysis, we scrutinized the isoform usage frequency variations across the periods surrounding ZGA, including those isoforms with very low expression but notable changes in usage. Such as *admp*, is known for its role in early dorsoventral patterning and was found to utilize a novel transcript pre-ZGA. Though expression of this gene is documented to initiate during the blastula stage, it is yet to be determined if the protein functionality is active, given that the transcript variant appears to lack coding capacity(*72*). Additionally, our findings indicate that the extent of transcript variation during ZGA is comparatively minimal, with most isoforms exhibiting differential usage without corresponding changes in gene expression. This suggests the crucial role of transcript stability in safeguarding developmental processes leading up to ZGA. Conversely, post-ZGA, transcript changes are more pronounced, with over half of these changes leading to differential gene expression. This indicates that ZGA activates the differential usage of a substantial number of transcripts, thereby driving the differential expression of related genes to facilitate rapid embryonic development. This underscores the critical role of ZGA in orchestrating the complex regulatory networks that drive early embryonic development. Moreover, transcript structure predictions suggest that ZGA preferentially utilizes transcripts with more complete functions, further underscoring the pivotal role of ZGA in early embryonic development. Unlike transcript usage, differential splicing events were already significantly occurring during ZGA. However, these differential splicing did not lead to a significant differential gene expression during ZGA, nor was there a necessary connection between changes in splicing events before and after ZGA and changes in gene expression, suggesting the distinct regulatory mechanisms governing gene expression and alternative splicing during early development.

More intriguingly, this study offers a novel perspective on the fascinating question of cell developmental fate in developmental biology by examining transcript-isoform and alternative splicing dynamics. Our findings reveal that cells can be clustered into two distinct subclusters based on transcript isoform diversity and coding capacities, with a clear chronological relationship in their subsequent developmental differentiation processes. Furthermore, our analysis of alternative splicing events and RBP activity indicates that the developmental trajectory characterized by faster differentiation progress exhibits more active splicing. This heightened splicing activity suggests a pivotal role for alternative splicing in driving the rapid differentiation of specific cell lineages. Based on these results, we infer that cell-type specific activity of isoform switching and splicing is a critical determinant in lineage differentiation. The dynamic regulation of transcript isoforms and splicing events not only contributes to the diversification of cellular functions but also orchestrates the precise timing of developmental transitions. This underscores the importance of transcriptomic diversity in shaping the developmental fate of cells and highlights the potential of targeting splicing mechanisms to influence cell differentiation pathways.

On the other hand, several limitations in this study need to be addressed in the future. The current single-cell long-read technology based on single-molecule sequencing is constrained by limited sequencing throughput, resulting in sparse data acquisition. Although applying analytical methods such as SEACell strategy can mitigate the disadvantages of data sparsity, and strict thresholds set in the analysis also ensure the reliability of the results, it is inevitable that some crucial transcripts may not be captured. In addition, the sequencing accuracy of single-molecule sequencing remains suboptimal, which can affect the precision of cell data segmentation in single-cell sequencing applications. Moreover, we encountered challenges in accurately distinguishing between zygotic and maternal transcripts. Consequently, we were unable to definitively ascertain whether the major contributions to transcript changes during development before and after ZGA were derived from maternal or zygotic genes. This limitation underscores the need for more refined techniques to differentiate between these transcript sources. Future efforts should focus on enhancing the throughput and accuracy of single-cell long-read sequencing technologies. Improvements in sequencing chemistry and bioinformatics algorithms are essential to achieve more comprehensive and precise transcriptome profiling(*73*). Additionally, developing better methods to distinguish between maternal and zygotic transcripts will be crucial for understanding the dynamics of gene regulation during early embryonic development. Furthermore, integrating multi-omics approaches, such as combining full-length transcriptomics with proteomics and epigenomics, will provide a more holistic view of the regulatory networks that drive cellular heterogeneity and development.

## Materials AND Methods

### Cell line and zebrafish embryos samples

Human embryonic kidney (HEK293T) and mouse fibroblast (NIH-3T3) cell lines, sourced from American Type Culture Collection (ATCC), were employed in this study. Additionally, we utilized AB wild-type zebrafish embryos, with all experiments approved by the Animal Care and Use Committee of Huazhong Agriculture University (HZAUFI-2021-0001). Embryos were harvested at various developmental stages post-fertilization: 128-cell (2.25 hpf), 512-cell (2.75 hpf), 1k-cell (3 hpf), Dome (4.3 hpf), and 50%-epiboly (5.25 hpf).

### Single cell suspension isolation and full-length cDNA construction

Single-cell suspensions from HEK293T and NIH-3T3 cell lines were prepared following the DNBelab C4 protocol (MGI, 940-001818-00). Zebrafish embryo single-cell suspensions were isolated using established methods(*26*). We processed 3,000 to 5,000 single cells using the DNBelab C-TaiM 4 (MGI, 900-000637-00) to generate barcoded single-cell cDNA libraries, adhering to the DNBelab C4 library construction protocol (MGI, 940-001818-00).

### Generating benchmarking data via parallel NGS of fragmented cDNA

A quarter (25% volume) of the barcoded full-length cDNA was subjected to fragmentation to prepare for short-read sequencing, adhering to the DNBelab C4 library construction protocol (MGI, 940-001818-00) and subsequently sequenced using a DNBSEQ-T20 sequencer. The raw data were processed using DNBC4tools (https://pypi.org/project/DNBC4tools/) to generate gene expression matrix.

### CycloneSEQ library preparation and sequencing

The remaining aliquot of the single-cell libraries underwent 15 cycles of amplification with cDNA Amp Primer-V3 and purified with 0.6x DNA Clean Beads from DNBelab C4 library construction kit (MGI, 900-000637-00). The integrity and size distribution of the full-length cDNA libraries were evaluated using an Agilent Bioanalyzer 2100 (Agilent Technologies, G2939A). Additionally, library purity was quantified by measuring the A260/A280 ratio with a NanoDrop spectrophotometer (Thermo Fisher Scientific, NanoDrop 2000). CycloneSEQ libraries were then constructed in accordance with the CycloneSEQ library kit instructions (CycloneSEQ, H940-000001-00) and subsequently sequenced using the CycloneSEQ WT02 platform.

### In situ hybridization

Custom primers, incorporating the T7 promoter sequence, were designed for the genes *wnt8a*, *myof* and *lsp1b*. The target fragments, generated through PCR amplification, were transcribed in vitro using T7 RNA Polymerase (Thermo Scientific, EP0111) and DIG RNA Labeling Mix (Sigma, 11277073910) to produce probes, which were stored at -80°C. Whole-mount in situ hybridization on 128-cell and shield-stage zebrafish embryos was conducted as previously described(*74*). Briefly, embryos were fixed in 4% PFA at 4°C, dehydrated in methanol, and stored at -20°C. After rehydration, they were washed in 1x PBST (Sigma, PPB005) and pre-hybridized at 70°C. Hybridization was performed with 1 ug/ml probe and RNase inhibitor (Takara, 2313A) at 70°C overnight. Post-hybridization washes included 2x SSC (Thermo Fisher Scientific, AM9770) at 70°C, followed by 0.2x SSC for 1 hour. Embryos were blocked with 1x PBST containing 2% goat serum and 2 mg/ml BSA (Thermo Fisher Scientific, 37525), then incubated with 1/5000 Anti-Digoxigenin-AP (Sigma, 11093274910) at 4°C. Final washes were in 1x PBST and Alkaline Tris buffer. Color development was achieved with NBT/BCIP Stock Solution (Sigma, 11681451001), followed by the application of a stop solution and storage or imaging at 4°C in the dark.

### Preprocessing of the CycloneSEQ read

Raw FASTQ files were preprocessed using Seqkit(*75*) to exclude reads shorter than 300 bp. Quality control was performed with Fastp(*76*), applying parameters -q 10 and -u 40. The quality-controlled reads were screened for expected adapters and filtered using Cutadapt to remove reads without adapters(*77*). Barcode demultiplexing was conducted using BWA by aligning the reads to a valid barcode list from DNBSEQ, with an allowed edit distance of 6 for 26-base pair barcodes(*78*). Subsequently, barcode reads were aligned to the zebrafish genome (GRCz11) using Minimap2(*79*) with the parameters (-ax splice, --secondary=no, -k14) to generate BAM files. UMI correction was then applied using UMI-tools(*80*), followed by quantification with Isoquant(*81*) to generate the single transcript matrix. Finally, the identified transcripts were classified and annotated using Sqanti3(*82*).

### Single cell short read data analysis

The single-cell gene expression matrix generated by DNBC4tools was processed for downstream analysis using the Seurat R package (v4.1.1)(*83*). Initially, we utilized DoubletFinder(*84*) to identify and remove potential doublets, and further filtered out cells with fewer than 500 genes expressed or with a mitochondrial percentage exceeding 5%. Following this, we conducted log normalization, proceeded with dimensionality reduction, and then performed clustering of cell types based on differential gene expression.

### Single cell long read data analysis

Single-cell long-read matrices obtained from Isoquant(*81*) were processed to filter out cells with transcript expression below 5 UMI and presence in at least 3 cells. To address sparsity in the transcript matrix, we employed SEACells to generate metacells(*54*). We then assigned cell annotation information from short-read data to long-read data based on cell IDs. Subsequently, we grouped cells into metacells within each cell type from the same developmental stage, ensuring that each metacell contained approximately 4,000 transcripts, based on expression similarity. Finally, we conducted downstream analysis using Scanpy(*85*), which involved log normalization and dimensionality reduction.

### Differential transcript usage (DTU) analysis

To assess differential transcript usage (DTU) between two consecutive developmental stages, we utilized IsoformSwitchAnalyzeR(*86*). The raw expression matrix for transcripts at each stage was randomly partitioned into two groups (each comprising 2 replicates), to generate bulk-level transcript and gene matrices. We then conducted DTU analysis between these stages using IsoformSwitchAnalyzeR, with a statistical cutoff set at p < 0.05 and a minimum absolute difference (|dif|) of greater than 0.1.

To elucidate differential transcript usage (DTU) at various stage or cluster levels, we performed single-cell DTU analysis. Initially, we constructed a transcript usage matrix from the metacell data, excluding transcripts expressed in fewer than 10% of cells across any stage or cluster. We then conducted two steps of differential testing: local and individual. Local differential testing identified transcripts with distinct usage patterns across at least two stages or clusters, using a significance threshold of p < 0.05 and |dif| > 0.1. Subsequently, individual differential testing was performed on the transcripts that met the criteria from the local testing to identify stage- and cluster-specific DTU, applying the same significance threshold of p < 0.05 and |dif| > 0.1.

### Differential percent spliced in (PSI) analysis

To evaluate differential PSI events across different stages or clusters, we merged the metacell transcript matrix with the splicing information generated by SUPPA2(*87*) to create a single-cell PSI matrix. For differential PSI analysis, each event was filtered to be expressed in at least 100 cells and in at least 10% of cells in any given stage or cluster. We conducted pairwise comparisons, ensuring a minimum of 50 effective cells in each group, and performed 100 permutation tests, following the methodology outlined in MARVEL(*88*). Significant differential splicing events were identified based on a threshold of p-value < 0.05 and |dpsi| > 0.05. For each event, modal identification can be performed using Anchor(*66*).

### RBP motif analysis

We utilized the rMAPS platform for the analysis of RNA-binding protein (RBP) motifs associated with alternative splicing events(*89*) (http://rmaps.cecsresearch.org). We initially downloaded 453 RNA-binding protein (RBP) binding sites from the oRNAment database(*90*)(https://rnabiology.ircm.qc.ca/oRNAment/downloads) and integrated them with the existing RBP motifs within the rMAPS platform. Subsequently, we input differential splicing events for predictive analysis, focusing on RBPs with homologs in zebrafish for further examination.

### Transcript sub-clustering analysis

We utilized ROGUE(*91*) to evaluate cellular heterogeneity at both the transcript and gene levels. We leveraged differentially expressed genes at the gene level, with a significance threshold of corrected p < 0.05 and |logFC| > 0.5, to perform isoform sub-clustering at a resolution of 1.6.

### Isoform trajectory analysis

We utilized the k-nearest neighbors (kNN) algorithm to perform trajectory analysis based on the isoform sub-cluster matrix. Briefly, we selected the 10 closest neighbors for temporal analysis between adjacent time points, requiring that at least 5 neighbors originated from the same cluster. Distances were measured to be less than 5 times the median total distance at that time point, using median absolute deviations (MADs) as the unit of measurement.

## ACKNOWLEDGMENTS

We thank all our teams’ members, the China National GeneBank (CNGB) and Zhejiang Key Laboratory of Spatial Omics for their support. Project analysis was performed on the STOmics Cloud (https://cloud.stomics.tech).

## Author Contributions

L.L., Z.D. and C.-Y. L. conceived the idea; C.-Y. L., C. L. and L.C. supervised the study; X.L., C. L. and C.-Y. L. designed the experiment; D.L. participated in the validation experiments with the help of M.H. and K.M.; K.Z., H.W., Y.W. and J.Z. collected the zebrafish embryo and generated single cell full-length cDNA libraries with the help of R.Z., G.H., Y.H. and J.P.; X.W. analyzed the data with the help of Z.Y., R.X., J.W., S.Y. and J.L.; T.Z., Y.D., J.C. and F.G. supported the single cell long read sequencing; X.L., C. L., L.C., X.W. and Z.Y. wrote the manuscript; L.L., Z.D., D.L., Y.G. and X.X. participated in the manuscript revised.

---

**fig. S1.**
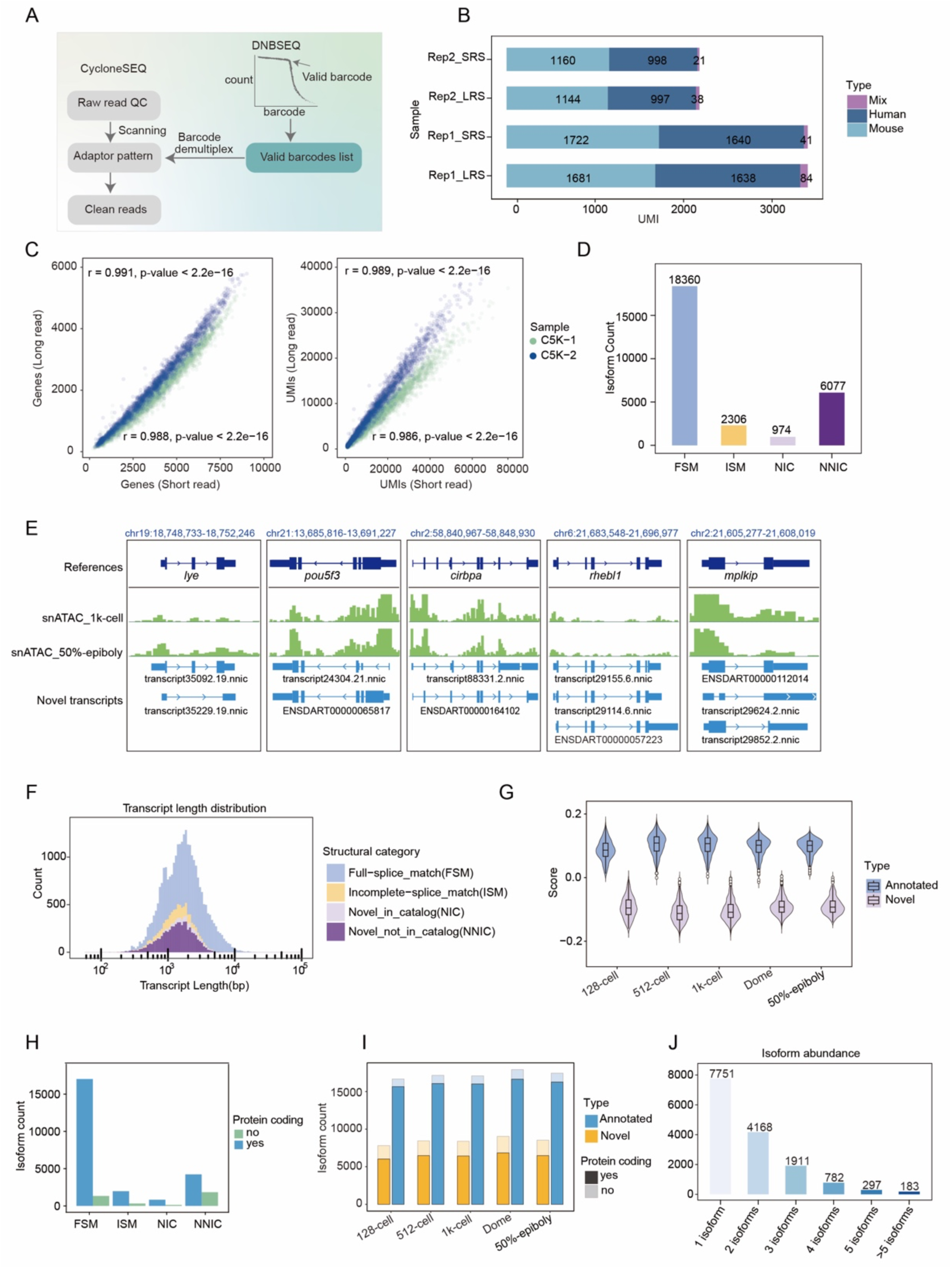
Full-length transcriptome diversity in zebrafish early embryogenesis. (A) Schematic diagram illustrating the long read single cell barcode demultiplexing. (B) The stacked histogram depicts UMI expression levels in mouse (NIH-3T) and human (HEK293T) cells across long read and short read data. (C) A scatter plot shows the correlation of gene and UMI counts per cell between short-read and long-read sequencing in C5K samples. (D) Bar chart showing the distribution of counts across various transcript structural categories. (E) Novel transcripts of the *lye*, *pou53*, *cirbpa*, *rhebl1* and *mplkip* genes were validated using snATAC-seq datasets, showing peak evidence at the transcription start site (TSS), as visualized in IGV. (F) A histogram displays the length distribution and counts of transcripts across different transcript structural categories. (G) A violin plot depicts the overall transcript expression scores for both annotated and novel transcripts across five developmental stages. (H) A bar chart illustrates the coding potential of transcripts, as predicted by CPAT, across different transcript structural categories. (I) A stacked bar chart depicts the coding potential of both annotated and novel transcripts across different developmental stages. (J) A bar chart shows the count of different isoform abundance per gene.

**fig. S2.**
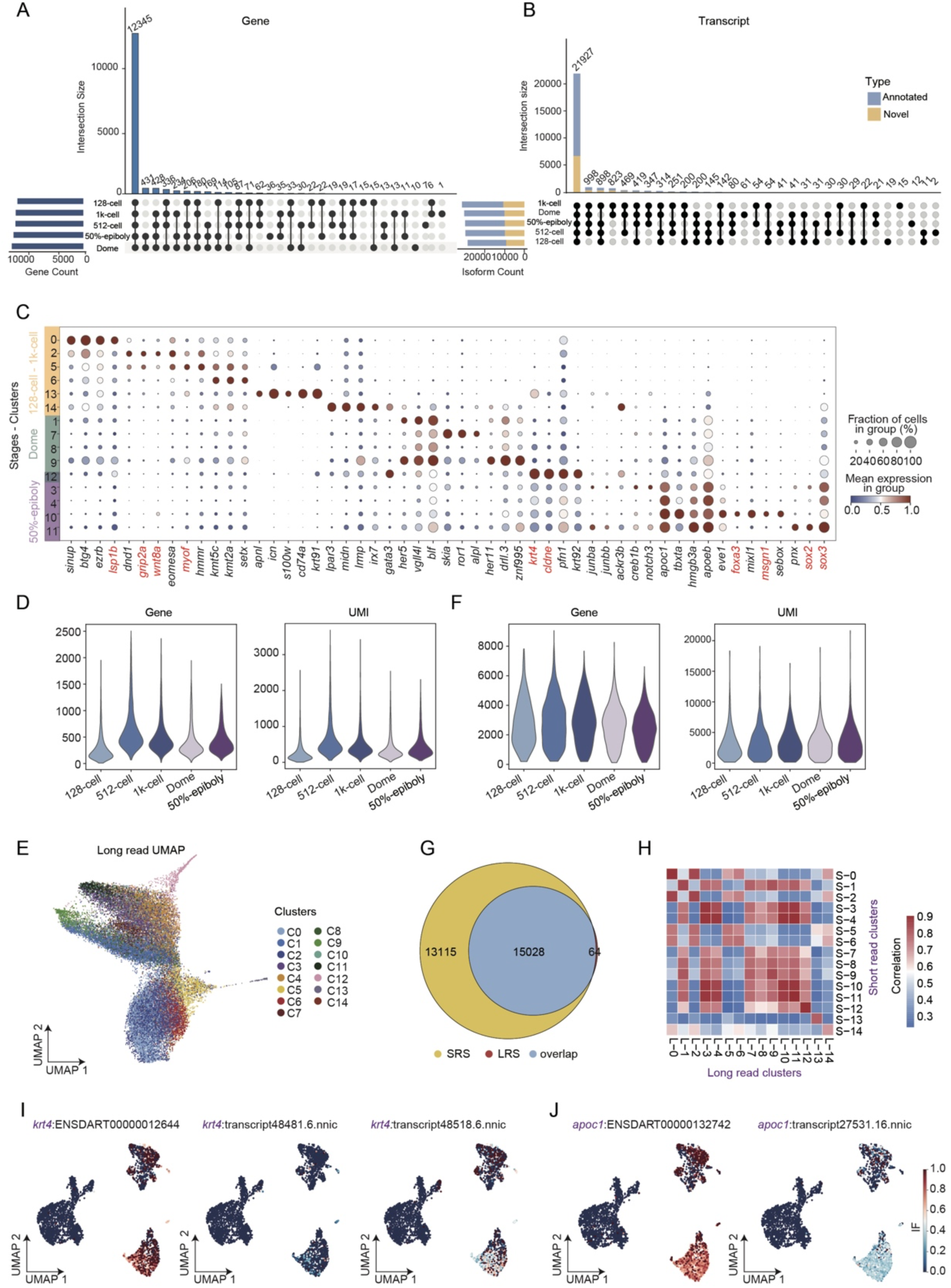
Full-length transcriptome diversity in zebrafish early embryogenesis. (A) An upset plot illustrates the counts of genes detected in long read data across five developmental stages. (B) An upset plot displays the counts of transcripts detected in long read data across five developmental stages. (C) A dotted heatmap illustrates the expression of cluster specific gene across cell clusters identified from short-read data. (D) A violin plot displays the distribution of genes and UMIs across different stages in long read data prior to the application of SEACells. (E) UMAP plot of 28,468 cells from long read sequencing data prior to the application of SEACells. (F) A violin plot displays the distribution of genes and UMIs across different stages in long read data following SEACells application. (G) Venn diagram showing the overlap of genes detected by long read and short read sequencing. (H) Heatmap showing the correlation of gene expression across cell clusters between long read and short read data. (I-J) A UMAP plot illustrates the distribution of isoform fractions (IF) among the transcripts of *krt4* (I) and *apoc1* (J) in long read data.

**fig. S3.**
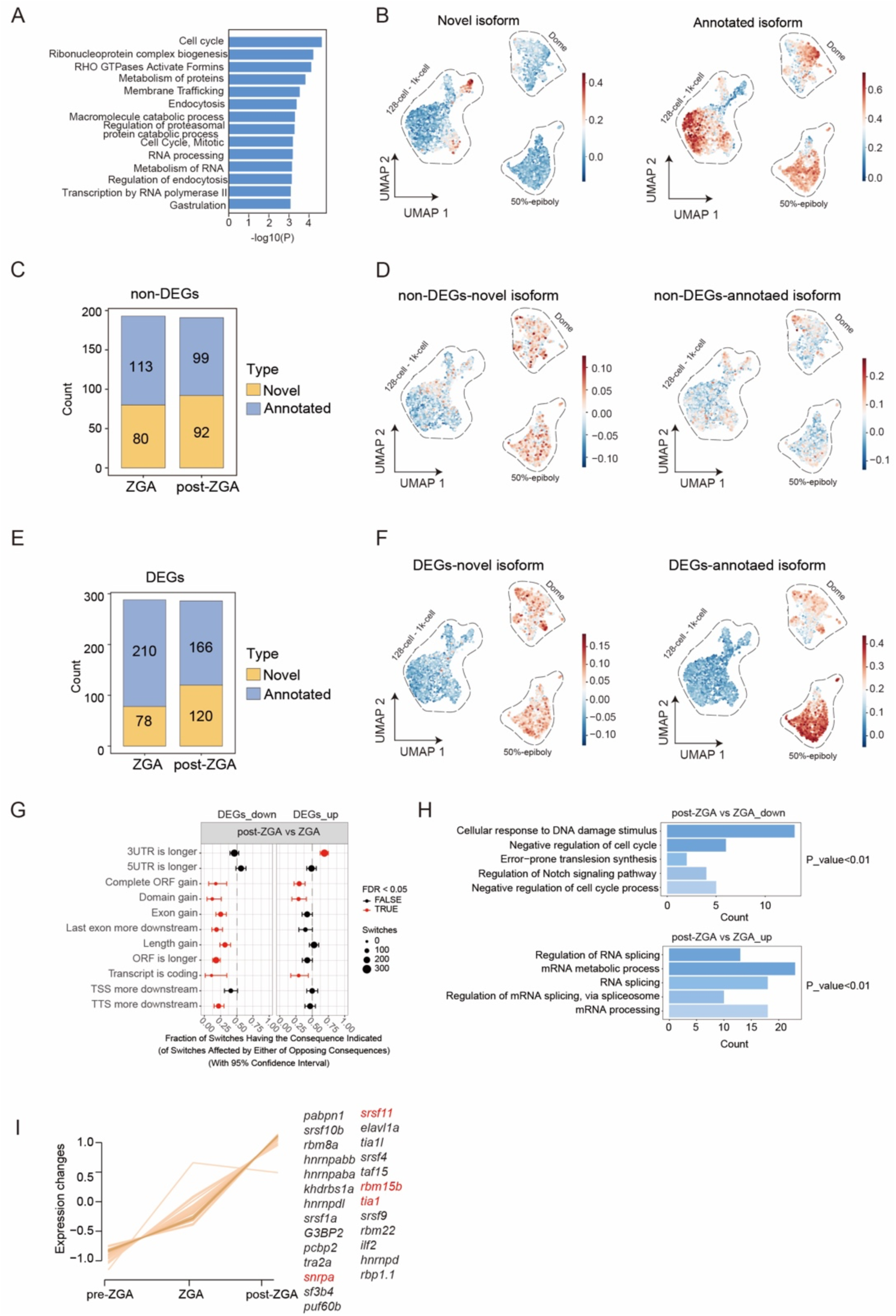
Global changes in transcriptome usage during zebrafish MZT. (A) A bar plot illustrates the GO enrichment analysis for genes associated with isoform switching in the non-DGE categories, comparing ZGA to pre-ZGA. (B) A UMAP plot displays the expression distribution of annotated and novel transcripts involved in isoform switching within the non-DGE categories, comparing ZGA to pre-ZGA. (C) The stacked bar chart displays the counts of annotated and novel transcripts, exhibiting isoform switching in non-DEG categories between ZGA and post-ZGA stages. (D) A UMAP plot displays the expression distribution of annotated and novel isoforms involved in isoform switching within the non-DGE categories, comparing post-ZGA to ZGA. (E) The stacked bar chart displays the counts of annotated and novel transcripts, exhibiting isoform switching in DEG categories between ZGA and post-ZGA stages. (F) A UMAP plot displays the expression distribution of annotated and novel isoforms involved in isoform switching within DGE categories, comparing post-ZGA to ZGA. (G) The illustration shows the functional consequences of isoform switching in DEGs categories (downregulated and upregulated), comparing post-ZGA to ZGA. (H) The bar graphs illustrate the GO enrichment analysis for genes with isoform switching, categorized by downregulation gene (top) and upregulation gene (bottom), comparing post-ZGA to ZGA. (I) The line graph showcases the expression levels of RNA-binding proteins (RBPs) that exhibit an increasing trend with stage progression.

**fig. S4.**
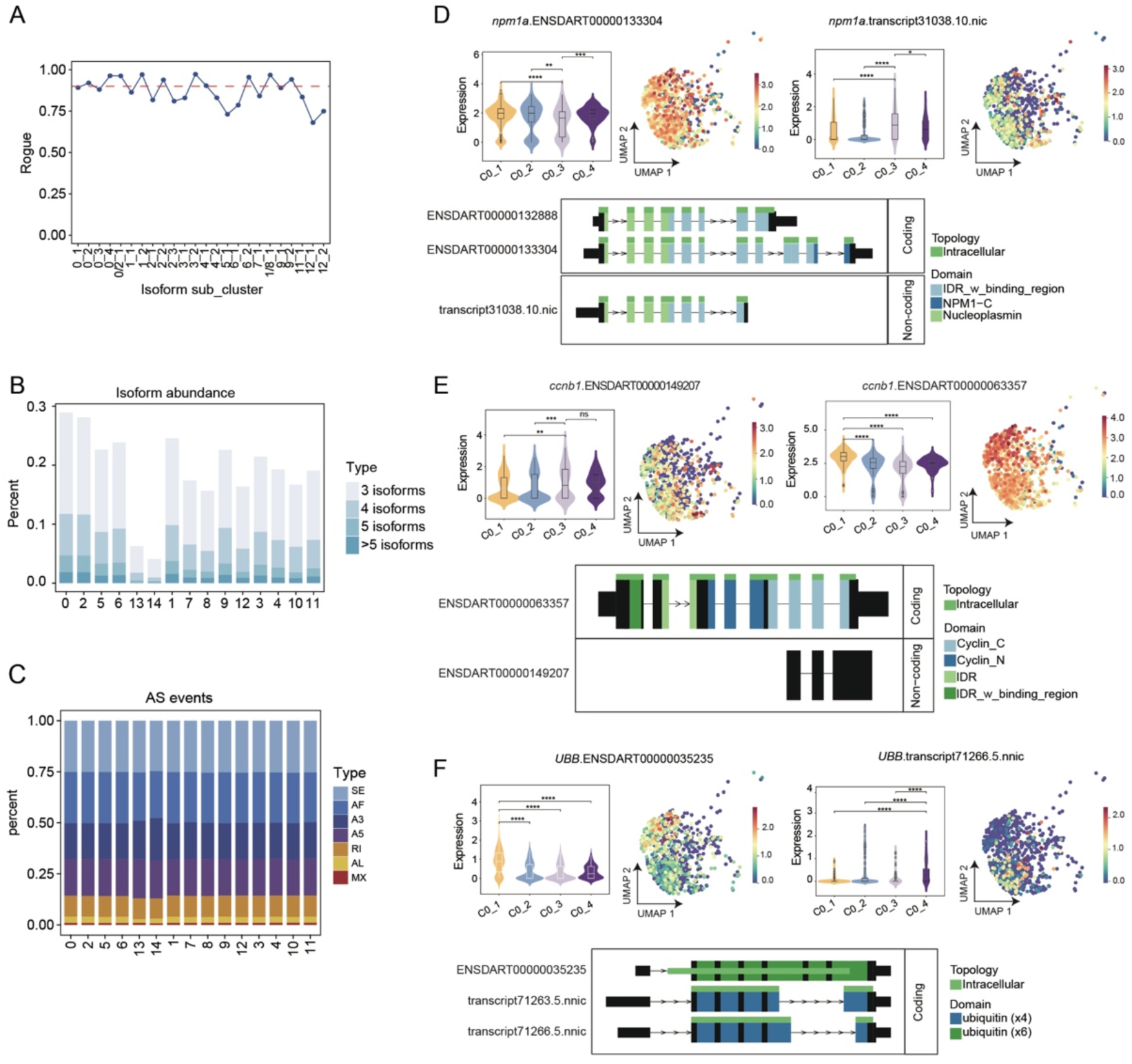
Isoform diversity uncovering finer subpopulations. (A) A line graph illustrates the assessment of cellular heterogeneity within the sub-clusters of long read transcript levels using ROGUE. (B) A stacked bar chart illustrates the transcript abundance per gene form long read data. (C) A stacked bar chart illustrates the detected splicing events in long read data, with clusters delineated at the gene level. (D-F) Violin and UMAP plots (top) display the expression distribution of different transcripts of *npm1a* (D), *ccb1* (E), and *ubb* (F) within the C0 sub-clusters. The switch plot (bottom) presents the results of transcript isoform switching related to the aforementioned genes.

**fig. S5.**
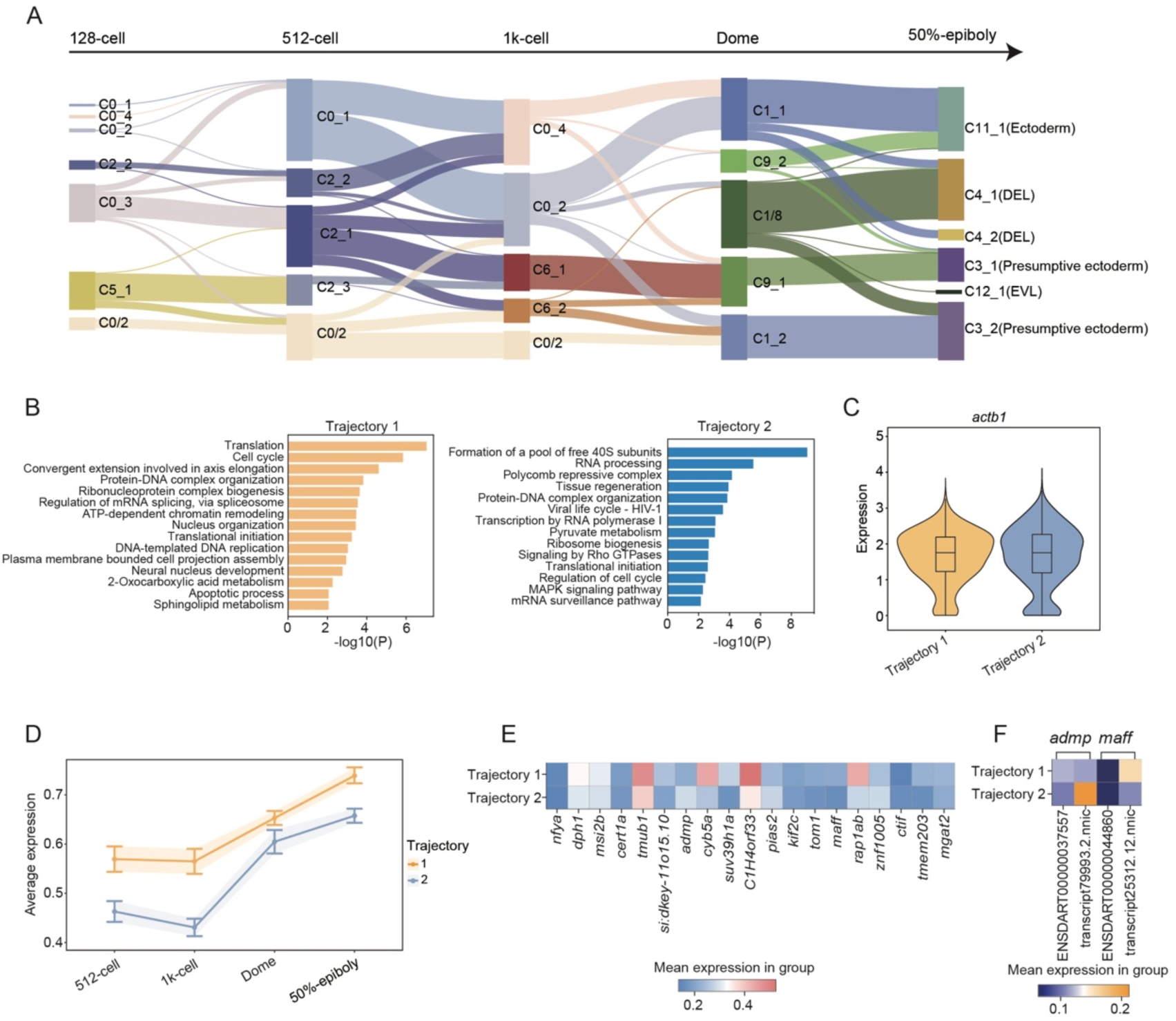
Isoform subgrouping associated with distinct developmental trajectories. (A) A Sankey diagram visualizes the developmental trajectory of zebrafish embryos, incorporating isoform sub-clusters identified in long read data. (B) A bar plot illustrates the GO enrichment of genes with differential exon usage between two trajectories with p < 0.05. (C) Violin plot showing the expression profile of the housekeeping gene actb1 across two developmental trajectories. (D) Expression dynamics of RBP genes identified via RNA-binding protein (RBP) motif prediction across two trajectories at different time points. (E) Heatmap of gene expression with significant differential exon usage between two trajectories. P < 0.05, ΔPSI > 0.3. (F) Heatmap displaying the expression patterns of distinct transcripts for *admp* and *maff* across two lineages.

